# Effects of Endogenous Angiotensin II on Abdominal Aortic Aneurysms and Atherosclerosis in Angiotensin II-infused Mice

**DOI:** 10.1101/2020.11.18.377416

**Authors:** Masayoshi Kukida, Hisashi Sawada, Satoko Ohno-Urabe, Deborah A. Howatt, Jessica J. Moorleghen, Marko Poglitsch, Alan Daugherty, Hong S. Lu

**Affiliations:** Saha Cardiovascular Research Center, Lexington, KY; Department of Physiology, University of Kentucky, Lexington, KY; Attoquant Diagnostics, Vienna, Austria

**Keywords:** angiotensin II, renin-angiotensin system, atherosclerosis, aortic aneurysms, endogenous

## Abstract

Angiotensin II (AngII), a major effector of the renin-angiotensin system, exerts critical roles in regulating vascular function. AngII infusion induces abdominal aortic aneurysms (AAAs) and exacerbates atherosclerosis in hypercholesterolemic mice. We determined the effects of AngII infusion on endogenous AngII regulation and AngII-mediated AAAs and atherosclerosis. AngII infusion increased renal, but not plasma, AngII concentrations in male mice. AngI concentrations were decreased modestly in kidney, but more profoundly in plasma, during AngII infusion. Bovine AngII (DRVYVHPF) has one amino acid difference from mouse AngII (DRVYIHPF) that can be distinguished by LC-MS/MS. Therefore, we determined exogenous versus endogenous peptides in mice infused with bovine AngII. Bovine AngII infusion reduced endogenous renal AngII concentrations. To determine whether the residual endogenous AngII exerted an effect on AAAs and atherosclerosis in mice infused with AngII, aliskiren (a direct renin inhibitor) was administered to AngII-infused male LDL receptor deficient mice. Although aliskiren did not attenuate AAAs in AngII-infused mice, atherosclerotic lesion size was reduced. In conclusion, endogenous AngII concentrations are reduced during AngII infusion but still contribute to atherosclerosis, but not AAA, in AngII-infused hypercholesterolemic mice.

---

Angiotensin II (AngII) is a major effector of the renin-angiotensin system and important in regulating vascular function. Infusion of AngII induces abdominal aortic aneurysms (AAAs) and exacerbates atherosclerosis in hypercholesterolemic mice. In AngII-infused normocholesterolemic rats, endogenous AngII production is maintained in the kidney.^1^ However, the effects of endogenous AngII on AAAs and atherosclerosis during AngII infusion in hypercholesterolemic mice have not been studied.

Our previous studies revealed that liver-specific deletion of angiotensinogen, the sole precursor of AngII, reduced atherosclerotic lesion area with decreases of AngII concentrations in kidney, but not in plasma.^2^ In addition, inhibition of angiotensinogen uptake into renal proximal tubular cells ameliorated atherosclerosis development.^3^ These results indicate an important role of renal AngII in atherosclerosis formation. In this study, either vehicle or murine AngII (1,000 ng/kg/min) was infused into male C57BL/6J mice for 7 days via osmotic pumps (Alzet model 2001, Durect Corporation, Cupertino, CA). Then, we determined concentrations of angiotensin peptides in plasma and kidney by liquid chromatography-mass spectrometry/mass spectroscopy (LS-MS/MS). Plasma AngII concentrations were not altered by AngII infusion (**Figure A**). Consistent with a previous report,^1^ renal AngII concentrations were significantly higher in AngII-infused mice than in vehicle-infused mice (**Figure A**). Since AngII is metabolized to AngIII and Ang(1-7) and these peptides may contribute to the pathophysiology of atherosclerosis,^4^ we assessed AngIII and Ang(1-7) concentrations in AngII-infused mice. Ang(1-7) was not detectable in plasma and kidney from either vehicle- or AngII-infused mice, and AngIII concentrations were not different in plasma and kidney between infusions (**Figure B**). These results indicate that exogenous AngII does not affect productions of major AngII metabolites.

**Figure.**
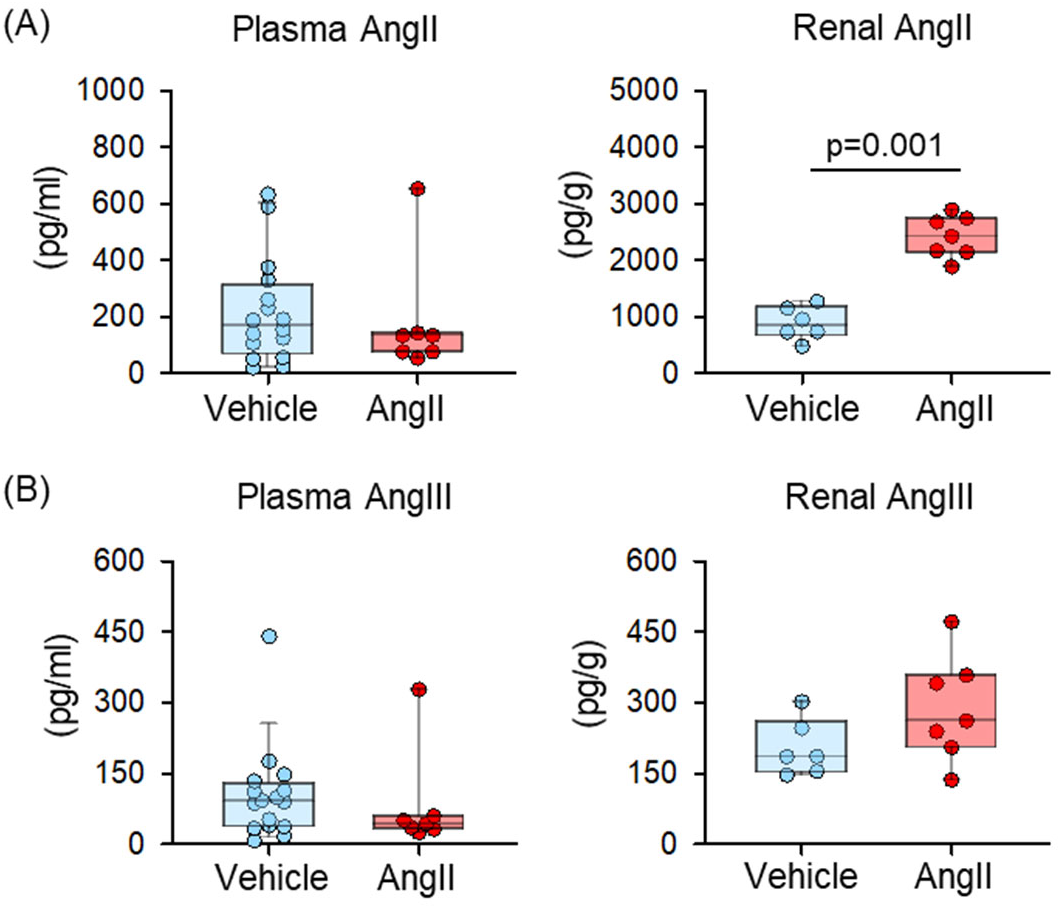
Plasma and renal concentrations of **(A)** Ang II and **(B)** AngIII in male C57BL/6J mice (8-10-week-old) infused with either vehicle (n=16) or murine AngII (1,000 ng/kg/min, n=7) for 7 days.

Since AngI is the direct substrate of AngII, we measured plasma and renal AngI concentrations to evaluate the production of AngII. AngII infusion decreased AngI concentrations in both plasma and kidney (**Figure C**). However, AngII-induced reduction of AngI concentrations was modest in kidney compared to those in plasma (96% in plasma vs 62 % in kidney, p<0.05 by Mann-Whitney U test, **Figure C**), indicating persistent renal AngII production during AngII infusion. We next investigated the presence of endogenous AngII in kidney of AngII-infused mice. Bovine AngII differs from murine AngII (Asp-Arg-Val-Tyr-**Ile**-His-Pro-Phe) in the fifth amino acid being Val (Asp-Arg-Val-Tyr-**Val**-His-Pro-Phe). The one amino acid difference results in a mass difference that enabled distinction between bovine and murine AngII by LC-MS/MS as exogenous and endogenous sources, respectively, in bovine AngII-infused mice. Bovine AngII exerted comparable effects to murine AngII on AAA and atherosclerosis formation in male LDL receptor deficient mice fed with Western diet (**Figure D**), indicating bovine AngII infusion mimicked murine AngII infusion-induced RAS regulation in mice. As expected, bovine AngII infusion increased exogenous AngII concentrations in kidney (**Figure E**). The median of renal endogenous AngII concentrations in vehicle-infused mice, was 843 pg/g (interquartile range: 669-1179 pg/g) (**Figure E**). AngII infusion reduced endogenous AngII concentrations to 161 pg/g (interquartile range: 77-235 pg/g). Endogenous AngII was still detectable in mice with bovine AngII infusion. Alongside the presence of renal AngI, these data support the notion that intrarenal AngII production is continued during AngII infusion.

**Figure (C).**
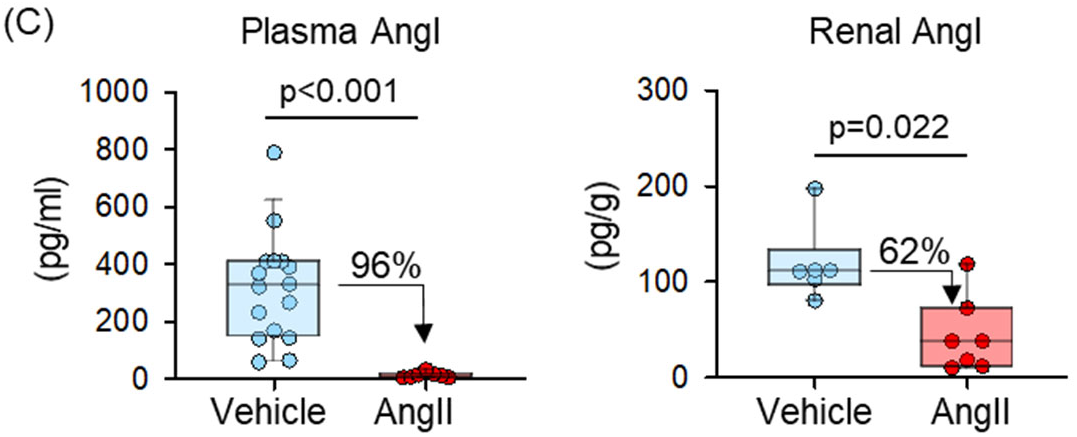
Plasma and renal AngI concentrations in male C57BL/6J mice infused with vehicle (n=16) or murine AngII (n=7) for 7 days.

**Figure (D).**
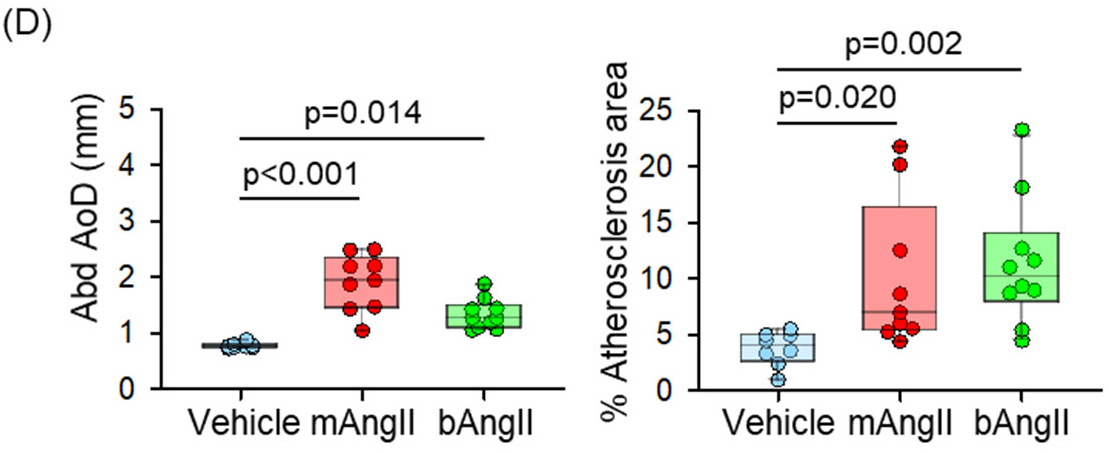
Maximal external diameters of abdominal aorta and percent atherosclerotic lesion areas in male LDL receptor deficient mice (10-week-old) fed Western diet and infused with vehicle (n=8), murine AngII (mAngII, 1,000 ng/kg/min, n=9), or bovine AngII (bAngII, 1,000 ng/kg/min, n=10).

**Figure (E).**
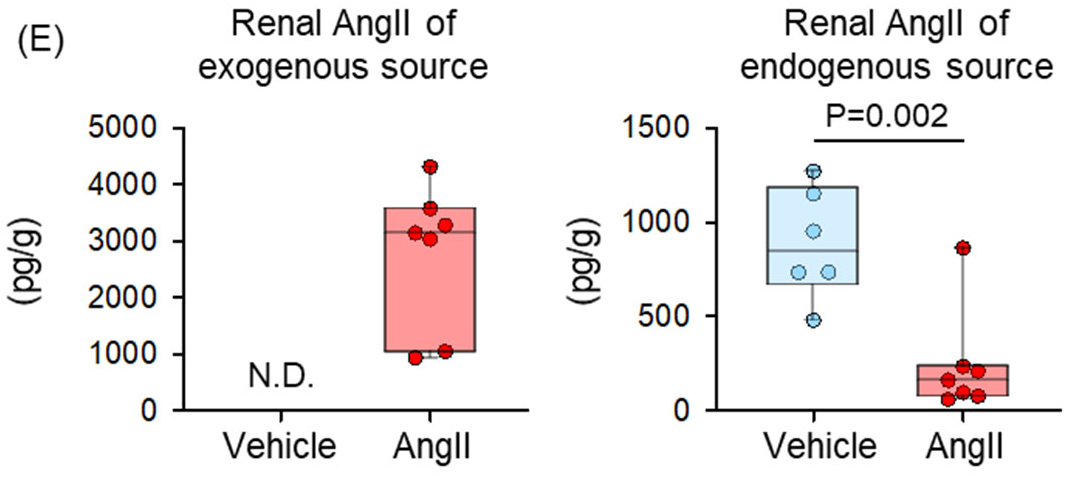
Renal AngII concentrations of exogenous and endogenous source in either vehicle (n=6) or bovine AngII (1,000 ng/kg/min, n=7) infused C57BL/6J mice.

We next investigated whether inhibition of endogenous AngII production attenuated AAA and atherosclerosis formation during AngII infusion. ACE inhibitors or direct renin inhibitors suppress endogenous AngII production. However, ACE inhibitors target several other substrates such as bradykinin that may affect cardiovascular functions. To investigate effects of endogenous AngII on AAA and atherosclerosis formation, the present study administered aliskiren, a direct renin inhibitor, to inhibit endogenous AngII production into AngII-infused mice. Murine AngII (1,000 ng/kg/min) was infused into LDL receptor deficient mice for 28 days that led to significant augmentation of AAA and atherosclerosis. Based on our previous study that aliskiren infusion of 25 mg/kg/day led to maximal inhibitory effects on endogenous AngII production in mice,^5^ we used this infusion rate of aliskiren in this experiment. As expected, continuous AngII infusion induced AAAs and augmented atherosclerotic lesions compared to vehicle infusion

(**Figure F**). Aliskiren did not inhibit abdominal aortic expansion in AngII-infused mice. Although atherosclerotic lesion size in aliskiren and AngII-infused mice was larger than vehicle-infused mice, aliskiren reduced lesion size significantly, compared to AngII infusion alone. These results support the concept that AAA development was attributed to exogenous AngII, whereas atherosclerosis was augmented by both endogenous and exogenous AngII.

**Figure (F).**
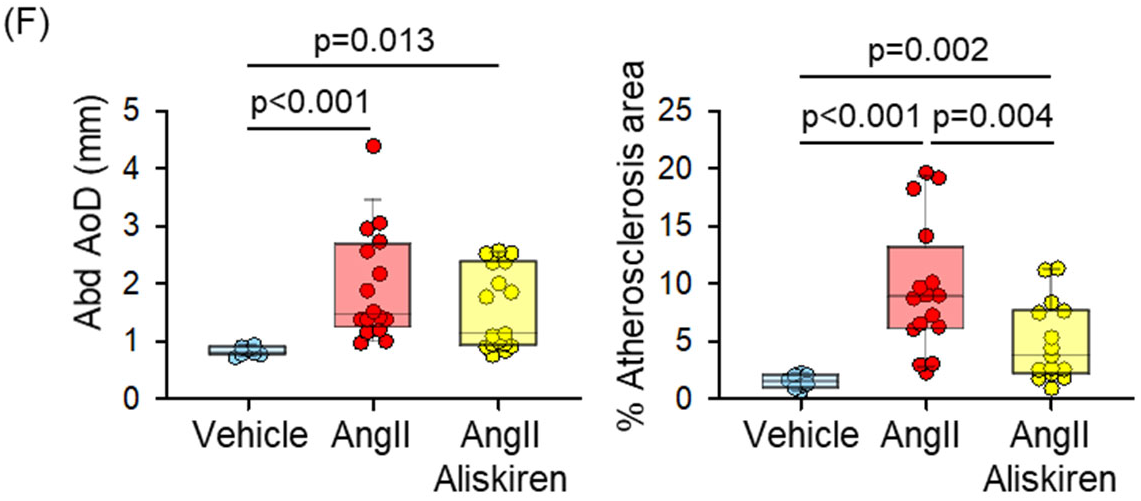
Aliskiren (25 mg/kg/day) suppressed atherosclerotic lesion size, but not abdominal aortic dilatation in LDLR deficient mice (8-week-old) with AngII infusion for 4 weeks (Vehicle;n=8, murine AngII;n=16, AngII+Aliskiren;n=17).

In conclusion, renal endogenous AngII is present in AngII-infused mice, and endogenous AngII contributes to AngII-mediated atherosclerosis, but not AAA, formation in hypercholesteremic mice.

## Future Perspective

While AngII infusion accelerates atherosclerosis formation and renal AngII concentrations, there has not been a determination that the renal changes directly impact the vascular pathology. Proximal tubule cells (PTCs) express all the components needed to generate AngII, including angiotensinogen, renin and ACE.^3^ AngII type 1a receptors, the determinant of AngII-induced atherosclerosis, are also present in PTCs. Therefore, we hypothesize that AngII production in PTCs augments atherosclerosis. To test this hypothesis, mice are developed with PTC-specific AngII type 1a receptor genetic deficiency or PTC overexpression of AGT and renin.

## Supporting information

raw data

## Abbreviations

AAA: Abdominal aortic aneurysm
ACE: Angiotensin-converting enzyme
Ang: Angiotensin
LS-MS/MS: Liquid chromatography-mass spectrometry/mass spectroscopy
PTC: Proximal tubule cell

## Acknowledgments

Study design: AD, HSL

Implementation of animal experiments: MK, HS, SO, JJM, DAH

Data analyses: MK, HS, HSL

LC-MS/MS and data interpretation: MP

Supervising and data verification: AD, HSL

Writing or editing the manuscript: MK, HS, HSL, AD

## Sources of Funding

This work was supported by NIH grants (R01HL139748, R01HL133723).

## Disclosures

None

## Methods

### Mice and Diet

C57BL/6J (Stock #000664) and low-density lipoprotein (LDL) receptor deficient (B6.129S7-Ldlr^tm1Her^; Stock #002207) male mice were purchased from The Jackson Laboratory (Bar Harbor, ME, USA). All mice were housed individually in ventilated cages, and fed a normal mouse laboratory diet. Mouse housing conditions are described in **Table I**. In experiments for atherosclerosis and AAAs, Western diet (**Table II**) was started one week prior to pump implantation and remained during AngII infusion. Because of the low incidence of AngII-induced AAAs in female mice, this study only used male mice.^6^ All experiments performed were approved by the University of Kentucky Institutional Animal Care and Use Committee.

**Table I.** Mouse Housing Conditions

| Condition | Mouse Housing Conditions | Note |
| --- | --- | --- |
| Cage Housing Intensity | 5 per cage |  |
| Light cycle (light:dark) | 14:10 hours |  |
| Water | Reverse Osmosis water system | ad libitum |
| Normal diet | Teklad 18% Protein Rodent Diet, # 2918, Envigo | ad libitum |
| Experimental diet for hypercholesteremia | Western diet, # TD.88137, Envigo | ad libitum for LDLR -/- mice |
| Bedding Source | Aspen hardwood chips, #7090A, Harlan Teklad Global |  |
| Temperature range | 18 – 23°C |  |
| Humidity range | 40 – 60 % |  |
| SPF | Yes |  |

**Table II.** Mouse Diet Information (Envigo)

|  | Normal diet<br>(weight/weight %) | Western diet<br>(weight/weight %) |
| --- | --- | --- |
| Diet # | 2918 | TD. 88137 |
| Protein | 18.6 | 17.3 |
| Carbohydrate | 44.2 | 48.5 |
| Fat | 6.2 | 21.2 |
| Cholesterol | 0 | 0.2 |

### Administration of AngII and aliskiren

To examine the profile of angiotensin peptides during AngII infusion, saline, murine AngII (mAngII, 1,000 ng/kg/min, Cat # H-1705, Bachem, Torrance, CA), or bovine AngII (bAngII, 1,000 ng/kg/min, Cat # H-1750, Bachem) was infused subcutaneously using osmotic mini pumps (Alzet model 2001, Durect Corporation, Cupertino, CA) for 7days into C57BL/6J mice at 8 −10 weeks of age, as described previously.^7^ Saline, mAngII, or bAngII was also infused into LDLR deficient mice at 10 weeks of age to compare effects between murine and bovine AngII on atherosclerosis and AAA formation.

For investigating the effects of aliskiren on AngII-mediated atherosclerosis and AAAs, either PBS or aliskiren (25 mg/kg/day, Novartis) was administered using another osmotic pump (Alzet model 2004) in addition to either saline or mAngII (1,000 ng/kg/min) infusion. Study groups with mouse numbers are described in the **Table III**.

**Table III.** Group designs in each figure

| Figure | Group |
| --- | --- |
| (A) - (C) | 1: vehicle (saline), n=16<br>2: murine AngII, n=7 |
| (D) | 1: vehicle (saline), n=8<br>2: murine AngII, n=9<br>3: bovine AngII, n=10 |
| (E) | 1: vehicle (saline), n=6<br>2: bovine AngII, n=7 |
| (F) | 1: vehicle (saline+PBS), n=8<br>2: murine AngII+PBS, n=16<br>3: murine AngII+Aliskiren, n=17 |

### Measurements of angiotensin peptides in plasma and kidney

Blood samples were collected into tubes with EDTA (final concentration: 1.8 mg/ml) by right ventricular punctures at termination. Approximately 300 μl blood was then put into tubes with a proteinase inhibitor cocktail (provided from Attoquant Diagnostics GmbH, Vienna, Austria) to prevent degradation of angiotensinogen and angiotensin peptides. The inhibitor cocktail contained broad spectrum inhibitors against metalloproteases (ethylenediaminetetraacetic acid, 1,10-phenanthroline), aspartic proteases (pepstatin A), cysteine proteases (p-hydroxymercuribenzoic acid), serine proteases (AEBSF) and specific inhibitors for renin and aminopeptidases A and N to a final concentration of 5% vol/vol. Following the immediate stabilization using the inhibitor cocktail, plasma was collected by centrifugation at 3,000 rcf for 10 minutes, 4°C, and frozen at −80°C until analysis.

After bleeding, the right atrium was incised, and cold saline (8 ml) was perfused through the left ventricle to remove blood from the systemic circulation. Following saline-perfusion, the whole right kidney was dissected out and immediately frozen in liquid nitrogen after removal of perirenal fat. For measurements of renal angiotensin peptides, the kidneys were homogenized using a pestle and mortar on liquid nitrogen. The frozen tissue powder was dissolved in a guanidine based denaturing extraction buffer on ice.

Thawed plasma and extracted renal samples were spiked with 200 pg stable isotope-labeled internal standards for individual angiotensin metabolites. Following C18-based solid-phase-extraction, samples were subjected to liquid chromatography-mass spectrometry/mass spectroscopy (LS-MS/MS) analysis using a reversed-phase analytical column (Acquity UPLC® C18, Waters) operating in line with a XEVO TQ-S triple quadrupole mass spectrometer (Waters) in MRM mode. Internal standards were used to correct for peptide recovery of the sample preparation procedure for each angiotensin metabolite in each individual sample. Concentrations of AngI, AngII, AngIII, Ang(1-7), and bovine AngII were calculated considering the corresponding response factors determined in appropriate calibration curves in original sample matrix, on condition that integrated signals exceeded a signal-to-noise ratio of 10.

### Quantification of abdominal aortic aneurysms and atherosclerosis

Abdominal aortic aneurysms (AAAs) were evaluated by *ex vivo* measurements of external aortic diameters, as described previously.^8, 9^ Briefly, the aorta was dissected from the ascending region to iliac bifurcation and fixed with neutrally buffered formalin (10% wt/vol) overnight at room temperature. Periaortic tissues were removed gently. Aortas were pinned and imaged with a mm ruler using a dissection scope (SMZ-800, Nikon, Tokyo) and a digital camera (DS-Ri1, Nikon). To quantify abdominal aortic dilation, maximal aortic diameters were measured at the most expanded region perpendicularly to the aortic axis using Image-Pro software (Media Cybernetics Inc., Rockville, MD). All measurements were verified by an investigator who was blinded to the study groups.

Atherosclerotic lesions were quantified on the intimal surface of aortic arches with an *en face* method following the AHA statement.^10^ After AAA measurements, the intimal surface was exposed by a longitudinal cut, and pinned on a black wax plate. Images of *en face* aortas were taken using the digital camera. Atherosclerotic measurements were performed using Image-Pro software. Lesion areas were traced manually in the ascending aorta, arch and part of the descending aorta (from the aortic orifice of left subclavian artery to 3 mm below). Lesion size was calculated as % lesion area using the following formula.

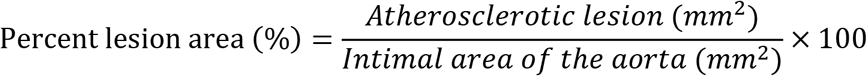

### Statistical analyses

SigmaPlot version 14.0 (Systat Software Inc., San Jose, CA) was used for statistical analyses. Normality and homogeneous variation were tested in all data by Shapiro-Wilk and Brown-Forsythe tests, respectively. Since the data did not pass either test, all data was analyzed using non-parametric analysis methods. For two-group comparisons, Mann-Whitney U test was performed. To compare multiple groups, Kruskal-Wallis oneway ANOVA on Ranks followed by Dunn’s method was used. All data was presented as box plots drawn from the 25th to 75th percentiles with a line at the median. P < 0.05 was considered statistically significant.

## Notes

### Competing Interest Statement

The authors have declared no competing interest.

### Summary of Updates

New data and detailed methods has been included in the revised version.

