## Supplementary material for "Effects of Endogenous Angiotensin II on Abdominal Aortic Aneurysms and Atherosclerosis in Angiotensin II-infused Mice": raw data

### Figure A

| Groups | Plasma AngII concentrations [pg/ml] |
|---|---|
| Vehicle-infused mice | 25 |
| | 56 |
| | 140 |
| | 19 |
| | 329 |
| | 187 |
| | 50 |
| | 106 |
| | 374 |
| | 588 |
| | 124 |
| | 633 |
| | 191 |
| | 261 |
| | 229 |
| | 154 |
| AngII-infused mice | 53 |
| | 133 |
| | 652 |
| | 142 |
| | 133 |
| | 76 |
| | 75 |

| Groups | Renal AngII concentrations [pg/g] |
|---|---|
| Vehicle-infused mice | 479 |
| | 735 |
| | 1149 |
| | 732 |
| | 951 |
| | 1269 |
| AngII-infused mice | 2140 |
| | 2166 |
| | 1885 |
| | 2677 |
| | 2739 |
| | 2419 |
| | 2888 |

**Figure B**

| Groups | Plasma AngIII concentrations [pg/ml] |
|---|---|
| Vehicle-infused mice | 17 |
| | 33 |
| | 89 |
| | 7 |
| | 98 |
| | 114 |
| | 39 |
| | 37 |
| | 176 |
| | 133 |
| | 52 |
| | 440 |
| | 86 |
| | 147 |
| | 112 |
| | 93 |
| AngII-infused mice | 35 |
| | 59 |
| | 328 |
| | 43 |
| | 50 |
| | 32 |
| | 24 |

| Groups | Renal AngIII concentrations [pg/g] |
|---|---|
| Vehicle-infused mice | 146 |
| | 302 |
| | 186 |
| | 185 |
| | 154 |
| | 246 |
| AngII-infused mice | 472 |
| | 136 |
| | 239 |
| | 358 |
| | 341 |
| | 205 |
| | 261 |

### Figure C

| Groups | Plasma AngI concentrations [pg/ml] |
|---|---|
| Vehicle-infused mice | 65 |
| | 143 |
| | 267 |
| | 59 |
| | 331 |
| | 369 |
| | 140 |
| | 410 |
| | 321 |
| | 789 |
| | 169 |
| | 411 |
| | 552 |
| | 390 |
| | 232 |
| | 409 |
| AngII-infused mice | 15 |
| | 33 |
| | 3 |
| | 8 |
| | 6 |
| | 18 |
| | 12 |

| Groups | Renal AngI concentrations [pg/g] |
|---|---|
| Vehicle-infused mice | 80 |
| | 197 |
| | 112 |
| | 111 |
| | 102 |
| | 112 |
| AngII-infused mice | 118 |
| | 6 |
| | 38 |
| | 5 |
| | 73 |
| | 38 |
| | 18 |

**Figure D**

| Groups | Maximal external diameter of abdomina aorta (mm) |
|---|---|
| Vehicle-infused mice | 0.73 |
| | 0.74 |
| | 0.75 |
| | 0.76 |
| | 0.77 |
| | 0.78 |
| | 0.80 |
| | 0.88 |
| mAngII-infused mice | 1.04 |
| | 1.43 |
| | 1.47 |
| | 1.87 |
| | 1.94 |
| | 2.19 |
| | 2.20 |
| | 2.48 |
| | 2.49 |
| bAngII-infused mice | 1.05 |
| | 1.07 |
| | 1.10 |
| | 1.19 |
| | 1.27 |
| | 1.27 |
| | 1.43 |
| | 1.44 |
| | 1.63 |
| | 1.88 |

| Groups | %Atherosclerosis area |
|---|---|
| Vehicle-infused mice | 0.97 |
| | 2.40 |
| | 3.29 |
| | 3.56 |
| | 4.47 |
| | 4.95 |
| | 4.97 |
| | 5.49 |
| mAngII-infused mice | 4.38 |
| | 5.23 |
| | 5.54 |
| | 5.98 |
| | 6.97 |
| | 8.62 |
| | 12.51 |
| | 20.16 |
| | 21.77 |
| bAngII-infused mice | 4.46 |
| | 5.42 |
| | 8.70 |
| | 8.96 |
| | 9.31 |
| | 11.01 |
| | 11.63 |
| | 12.67 |
| | 18.13 |
| | 23.25 |

**Figure E**

| Groups | Renal AngII of exogenous source [pg/g] |
|---|---|
| Vehicle-infused mice | N.D. |
| | N.D. |
| | N.D. |
| | N.D. |
| | N.D. |
| | N.D. |
| AngII-infused mice | 3139 |
| | 1045 |
| | 3033 |
| | 934 |
| | 4309 |
| | 3276 |
| | 3572 |

| Groups | Renal AngII of endogenous source [pg/g] |
|---|---|
| Vehicle-infused mice | 479 |
| | 735 |
| | 1149 |
| | 732 |
| | 951 |
| | 1269 |
| AngII-infused mice | 77 |
| | 235 |
| | 60 |
| | 863 |
| | 96 |
| | 210 |
| | 161 |

**Figure F**

| Groups | Maximal external diameter of abdomina aorta (mm) |
|---|---|
| Vehicle-infused mice | 0.93 |
| | 0.90 |
| | 0.72 |
| | 0.77 |
| | 0.79 |
| | 0.86 |
| | 0.78 |
| AngII-infused mice | 0.97 |
| | 1.00 |
| | 1.16 |
| | 1.19 |
| | 1.38 |
| | 1.38 |
| | 1.38 |
| | 1.42 |
| | 1.51 |
| | 1.88 |
| | 2.17 |
| | 2.56 |
| | 2.73 |
| | 2.95 |
| | 3.05 |
| | 4.39 |
| AngII- and Aliskiren-infused mice | 0.76 |
| | 0.83 |
| | 0.90 |
| | 0.91 |
| | 0.94 |
| | 0.96 |
| | 0.99 |
| | 1.09 |
| | 1.13 |
| | 1.77 |
| | 1.85 |
| | 2.00 |
| | 2.37 |
| | 2.38 |
| | 2.52 |
| | 2.53 |
| | 2.57 |

| Groups | %Atherosclerosis area |
|---|---|
| Vehicle-infused mice | 0.71 |
| | 1.63 |
| | 2.23 |
| | 2.02 |
| | 2.06 |
| | 1.11 |
| | 1.30 |
| | 0.92 |
| AngII-infused mice | 2.28 |
| | 2.96 |
| | 3.05 |
| | 6.05 |
| | 6.24 |
| | 6.51 |
| | 7.24 |
| | 8.72 |
| | 8.96 |
| | 9.01 |
| | 9.67 |
| | 10.10 |
| | 14.13 |
| | 18.23 |
| | 19.17 |
| | 19.63 |
| AngII- and Aliskiren-infused mice | 0.91 |
| | 1.76 |
| | 1.82 |
| | 2.16 |
| | 2.53 |
| | 2.56 |
| | 2.65 |
| | 3.73 |
| | 4.42 |
| | 5.32 |
| | 7.47 |
| | 7.62 |
| | 8.36 |
| | 11.18 |
| | 11.32 |
